# Identification and removal of sequencing artifacts produced by mispriming during reverse transcription in multiple RNA-seq technologies

**DOI:** 10.1101/339887

**Authors:** Haridha Shivram, Vishwanath R. Iyer

**Affiliations:** Center for Systems and Synthetic Biology, Institute for Cellular and Molecular Biology, Department of Molecular Biosciences, University of Texas at Austin, Austin, Texas 78712, United States of America

**Keywords:** RNA sequencing (RNA-seq), Reverse transcriptase, Reverse transcription, Mispriming, Artifacts, EZH2, Polycomb repressive complex (PRC2), RNA-binding, GRO-seq, HITS-CLIP, Short RNA-seq, TGIRT

## Abstract

The quality of RNA sequencing data relies on specific priming by the primer used for reverse transcription (RT-primer). Non-specific annealing of the RT-primer to the RNA template can generate reads with incorrect cDNA ends and can cause misinterpretation of data (RT mispriming). This kind of artifact in RNA-seq based technologies is underappreciated and currently no adequate tools exist to computationally remove them from published datasets. We show that mispriming can occur with as little as 2 bases of complementarity at the 3’ end of the primer followed by intermittent regions of complementarity. We also provide a computational pipeline that identifies cDNA reads produced from RT mispriming, allowing users to filter them out from any aligned dataset. Using this analysis pipeline, we identify thousands of mispriming events in a dozen published datasets from diverse technologies including short RNA-seq, total/mRNA-seq, HITS-CLIP and GRO-seq. We further show how RT-mispriming can lead to misinterpretation of data. In addition to providing a solution to computationally remove RT-misprimed reads, we also propose an experimental solution to avoid RT-mispriming by performing RNA-seq using thermostable group II intron derived reverse transcriptase (TGIRT-seq).

## Introduction

RNA-seq technologies are widely used to address biological questions relevant to transcriptional, co-transcriptional and post-transcriptional regulation of gene expression. Some methods involve measurement of read coverage across an entire gene or exon while others utilize the specific positions of read pile-ups. A key step in all these RNA-seq technologies involves reverse transcription followed by library construction and sequencing. In some experiments, RNA adapters are first ligated to RNA 3’ ends followed by reverse transcription (RT) using a primer complementary to the ligated adapter (RT-primer). Alternatively, RT is first performed using random primers and then adapters are ligated to the cDNA molecules (Fig. 1A). The latter approach is utilized by standard Illumina TruSeq RNA-seq kits. The former is a cost-efficient approach to retain strand information and is thus used in many technologies [1–4]. Accurate cDNA synthesis relies on binding of the RT-primer specifically to the 3’ adapter ligated to RNA or the unbiased pairing of random primer to RNA. Non-specific binding of the RT-primer can produce artefactual reads due to RT mispriming and lead to misinterpretations of read counts and cDNA lengths in RNA-seq experiments (Fig. 1A) [5].

**Figure 1.**
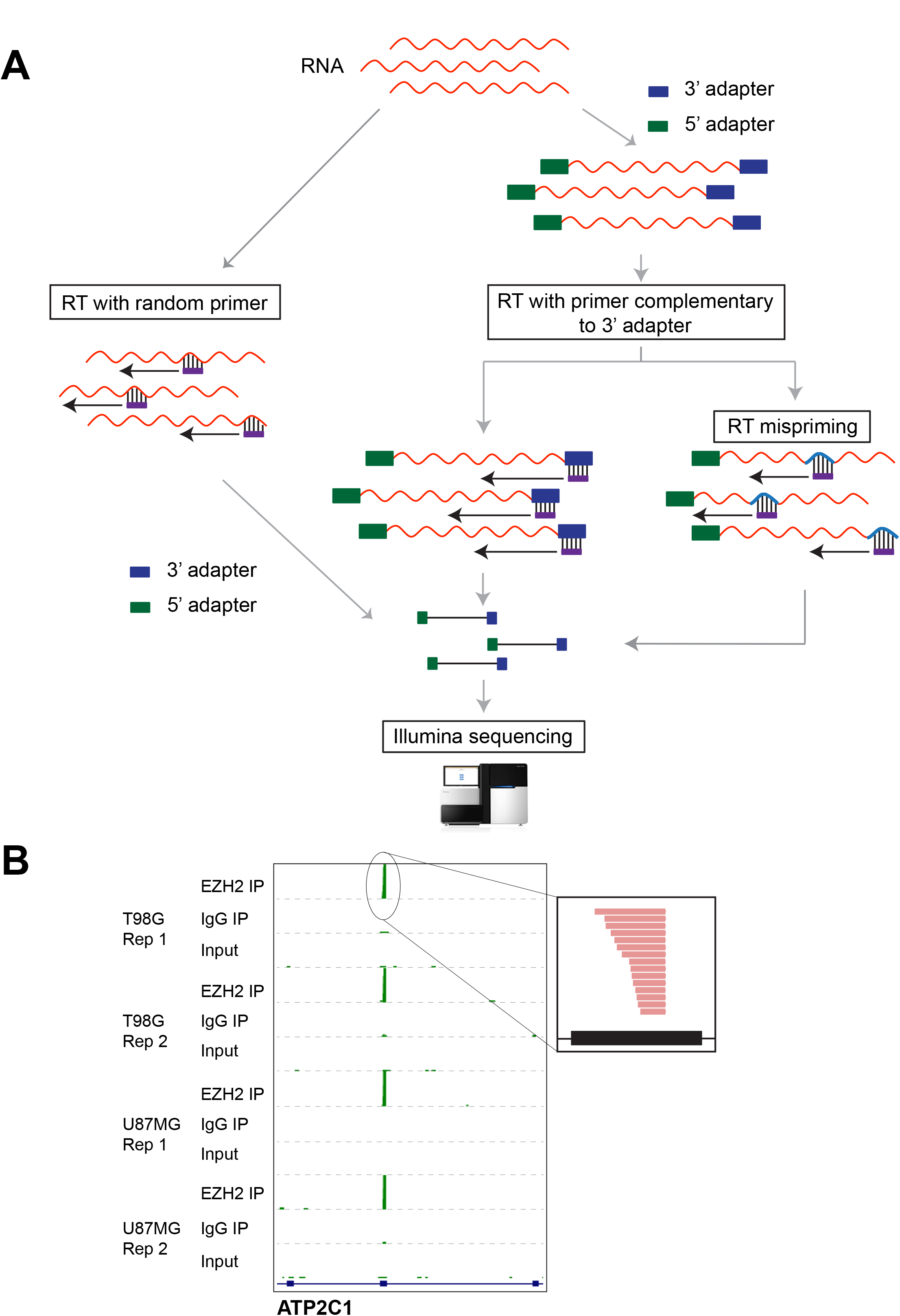
(A) Approaches for making cDNA libraries. One approach involves reverse transcription with random primers first, followed by adapter ligations and sequencing (left). The other approach is to first sequentially ligate 3′ and 5′ adapters, then perform cDNA synthesis using a primer complementary to the adapter (RT-primer) followed by sequencing (right). On using RT-primer with a specific sequence, mispriming could occur due to annealing of the RT-primer to transcript sequences with some complementarity (RT mispriming). (B) Genome browser view showing enrichment of sequencing reads at a specific exon in EZH2 short RNA RIP compared to IgG IP and input controls across different cell lines and independent replicates (Rep). The inset shows sense-strand reads (red) mapping to the peak region.

Aside from a couple of publications, RT mispriming has not been recognized as a potential problem in transcriptome-wide studies. Experimental methods to avoid RT mispriming artifacts were recently proposed for mammalian NET-seq (Native elongating transcript sequencing) and HITS-CLIP (High throughput sequencing of RNA isolated by crosslinking and immunoprecipitation) [6–9]. Although these methods will be useful for future experiments, there is still a need to identify and remove misprimed reads from existing datasets. Failure to account for or remove reads produced from mispriming during analysis of published datasets can lead to misinterpretation of data.

The current approach to identifying RT mispriming events involves looking for genomic regions close to cDNA peaks that are complementary to the first 6-7 bases of the RT-primer (matching the 3’ adapter) [6,7]. We however find that mispriming can occur with just 2 bases followed by scattered complementarity to the RT-primer. Thus, existing approaches underestimate the extent of mispriming in the data. Here, we provide an analysis pipeline to remove RT-misprimed reads and apply this to several published datasets. Using this approach we identify RT mispriming events in data from multiple RNA-seq technologies including HITS-CLIP, short RNA-seq, total/mRNA-seq and GRO-seq (Global run-on followed by sequencing) and further show how RT mispriming could lead to misinterpretation of data [10]. As an alternative to existing solutions, we propose cDNA library construction using the template-switching activity of novel thermostable group II intron-encoded reverse transcriptases (TGIRT-seq) as a reliable approach to avoid RT mispriming [11,12].

## Results

### Short RNA sequencing experiments show spurious peaks from coding exons

To identify short RNAs interacting with a chromatin modifier protein, EZH2, we performed a modified RIP-seq approach where we omitted all RNA digestion steps but instead size selected for 20-50 nt long RNAs [13]. We performed replicate short RNA RIP-seq experiments for EZH2 and analyzed them by comparing to two negative controls – an immunoprecipitation (IP) with non-specific IgG and input RNA that was not subject to IP but otherwise processed in parallel – in two glioblastoma multiforme (GBM) cell lines. We found RNA sense-strand reads piling up as peaks localized to a short region within specific exons from several genes (Fig 1B). Although we found several hundreds of these exonic cDNA peaks to be highly enriched in the EZH2 IP compared to the IgG control IP (fold change > 2 and FDR-corrected *P*-value < 0.05 using DESeq2), these peaks were detectable in both controls (Figure S1A).

Strikingly, the exonic cDNA peaks showed flush 3’ ends suggestive of mis-alignment or spliced reads (Fig 1B, Fig. 2A and Figure S1A-C). The raw reads from these cDNA peaks however showed no evidence of a novel splice junction or misalignment. To further understand their identity we checked for sequence biases at genomic regions flanking the short RNA peaks. We found sequences similar to the first few bases of the 3’ adapter adjacent to the short RNA peaks (Figure S1B, C). The nucleotide composition at genomic regions adjacent to the 3’ ends of all EZH2-enriched short RNA peaks showed a clear bias for a sequence similar to the 3’ adapter (Fig. 2B). Recently, similar peaks enriched for the 3’ adapter sequence were identified in HITS-CLIP data that were tagged as false positive peaks produced by mispriming during reverse transcription [7].

**Figure 2.**
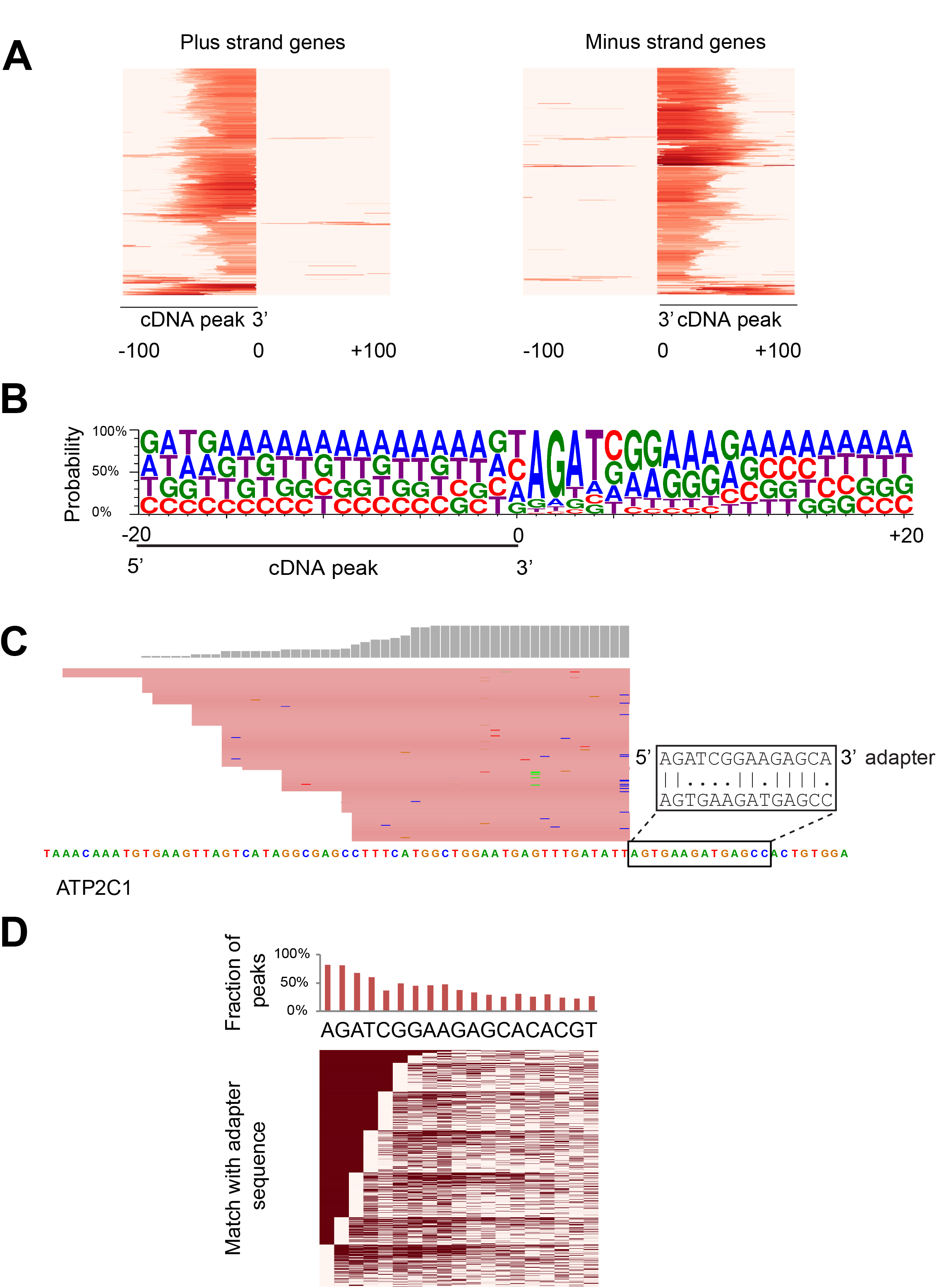
(A) Heatmap showing EZH2-enriched cDNA peaks with flush ends. The heatmap shows the distribution of reads spanning 200 bases around EZH2-enriched peaks. The intensity of red color represents mapped read counts. Each row represents an EZH2-enriched cDNA peak. (B) Sequence logo showing sequence enrichment spanning 40 bp around an EZH2-enriched cDNA peak with its 3′ end at position 0. (C) A cDNA peak adjacent to a sequence with its first 2 bases matching the 3′ adapter followed by scattered matches. (D) Sequence matches between bases adjacent to the 3′ end of all EZH2-enriched cDNA peaks and the sequence of the 3′ adapter (showing up to 19 bases). Top: Overall proportion of peaks showing a match at each position. Bottom: Heatmap of individual matches. Filled cells represent a match and empty cells correspond to positions that do not match the 3’ adapter. Rows are ordered by the number of matches starting from the left, which corresponds to the 3′ end of the adapter.

Mispriming sites were previously identified by looking for regions matching the first 6-7 bases of the 3’ adapter proximal to cDNA peaks. We however found artefactual exonic cDNA peaks produced from genomic regions with only partial matches to the first 6-7 bases of the 3’ adapter. For 80% of all EZH2-enriched exonic cDNA peaks, only the first two bases matched the 3’ adapter, and for 48%, the first seven bases matched the 3’ adapter with two mismatches allowed (Fig. 2C, D and Figure S1C). This suggests that based on the criteria previously used (7 bases complementary to the RT-primer with 2 mismatches allowed), only 48% of the exonic cDNA peaks would be identified as a false positive mispriming artifact.

### Pipeline to identify sites of mispriming from RNA sequencing datasets

The short RNA library preparation protocol involves ligation of the 3’ adapter followed by reverse transcription with an RT-primer complementary to the adapter [4,14]. False cDNA peaks are produced when the RT-primer binds to regions of complementarity on the RNA molecule and synthesizes cDNA (Fig. 3A). Based on the properties we observed for cDNA peaks in short RNA-seq experiments, we defined the following criteria to identify mispriming artifact peaks as distinct from true cDNA peaks (Fig. 3B). (1) Sites of mispriming should have at least two bases matching the 3’ end of the 3’ adapter, (2) cDNA peaks produced from mispriming should have flush 3’ ends with at least 10 reads high pile-up and, (3) There should be no other cDNA peak with 3’ flush ends resembling the misprimed peak but not matching the RT-primer within 20 bases flanking the misprimed peak. This is critical to avoid false mispriming calls at regions of high expression that likely contain many reads with flush ends as a result of high read density. We implemented these criteria in a computational pipeline to identify mispriming sites. The first step in the pipeline is alignment of sequencing reads using a global aligner (BWA). Since miRNAs and other small RNAs have defined ends, we then filter reads that do not map to nonprotein-coding genes. From the filtered alignment file, we identify genomic positions where cDNA peaks with flush ends (> 10 reads) are adjacent to (1) dinucleotides matching the 3’ adapter (k-mer sites) and, (2) dinucleotides that do not match the 3’ adapter (non-k-mer sites). Finally, mispriming sites are identified as k-mer-sites that do not contain a non-k-mer site within 20 bases. For datasets containing misprimed reads, a significant fraction of mispriming sites identified by our pipeline is expected to match more than the first two bases of the 3’ adapter.

**Figure 3.**
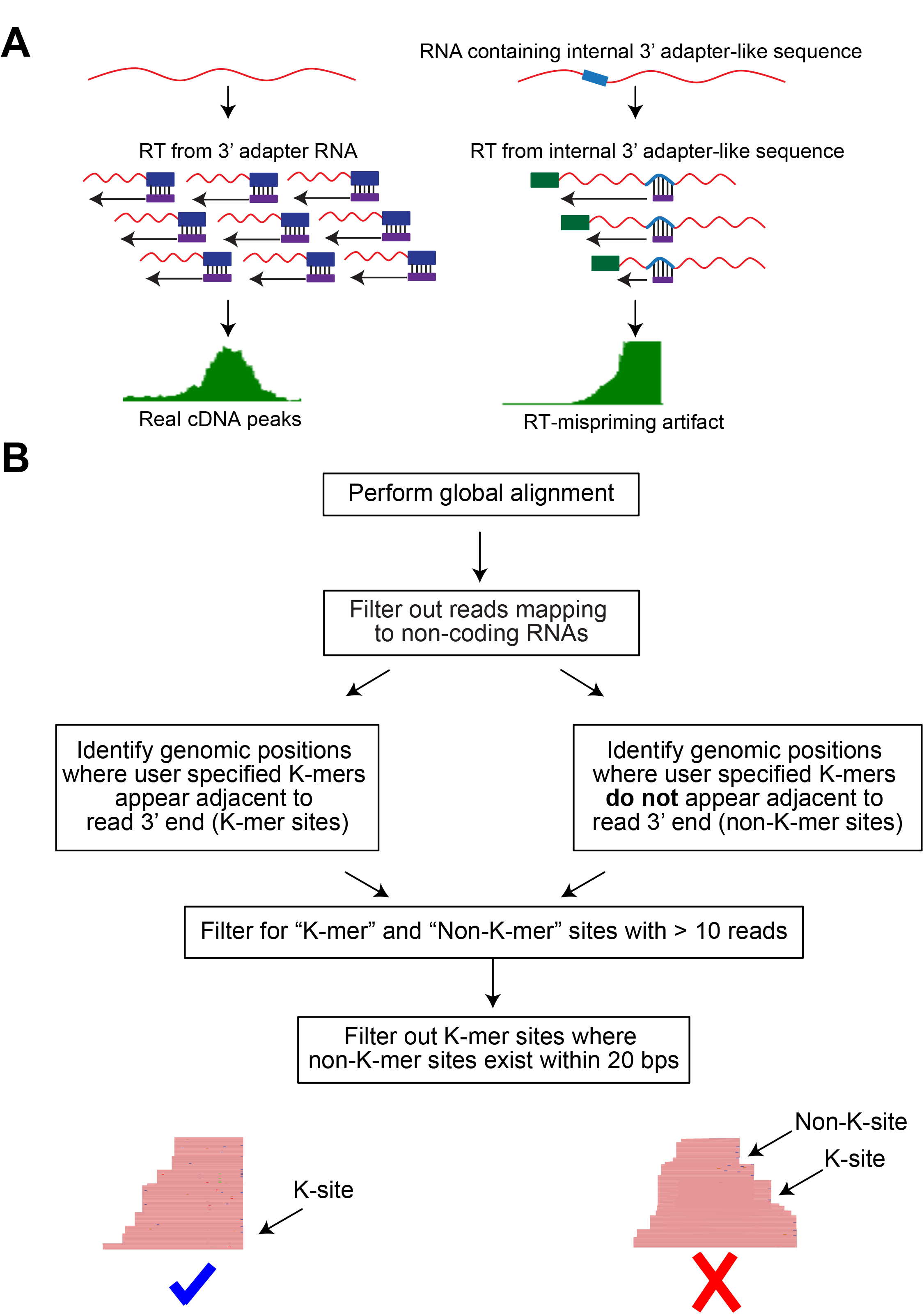
(A) Schematic comparing bonafide cDNA peaks with peaks from mispriming events. RNA molecules that are properly ligated and reverse transcribed from specific RT primer-3’ adapter interaction produce a pile-up of cDNA reads that have staggered ends (left). On the other hand, when RT-primer pairs with a sequence similar to the 3’ adapter present within an RNA molecule, cDNA peaks with flush ends next to the priming site are produced. (B) Pipeline to identify sites of mispriming.

With this approach we were able to identify mispriming sites in several short RNA-seq datasets with ~95% success rate (~1700 out of ~1800 short RNA peaks containing 2 bases complementary to the RT-primer). As expected, we found enrichment for the 3’ adapter sequence downstream of mispriming sites identified by our pipeline in all short RNA-seq datasets (Figure S2A). By identifying precisely the sites of mispriming, we were also able to filter out reads that were produced as a result of RT mispriming (Figure S2B). In addition to short RNA peaks enriched in EZH2 IP samples, we were able to identify more than 10,000 mispriming sites per dataset (Table 1). We found that the number of mispriming sites decreased as the complexity of the library increased, with the input library showing the least amount of mispriming. We also checked for mispriming in an independent short RNA-seq dataset that we downloaded from GEO (GSE68254) [15]. Similar to our input short RNA-seq samples, we also observed thousands of mispriming sites in this published dataset (Table 2, Figure S2C). Since we were able to detect mispriming events in short RNA-seq input libraries, we suspected that mRNA-seq libraries might also be contaminated with misprimed reads. Using our pipeline, we identified several misprimed reads in total/mRNA-seq datasets we generated (T98G and U87MG cells) that coincided with the position of misprimed peaks from short RNA-seq experiments. Similarly, we also found several mispriming events in published mRNA-seq datasets that we downloaded from GEO (Figure S3).

**Table 1:** RT mispriming events identified by our pipeline in published datasets.

| Study | Accession | Technique | Number of mispriming sites | Number of misprimed reads | Percent of reads misprimed |
| --- | --- | --- | --- | --- | --- |
| GSE68254 | SRR1995349 | Short RNA-seq | 2773 | 75105 | 0.587 |
| GSE63556 | SRR1660042 | RNA-seq | 4004 | 55688 | 0.082 |
| GSE63556 | SRR1660043-45 | RNA-seq | 3858 | 44165 | 0.073 |
| GSE63496 | SRR1658024 | RNA-seq | 122 | 3145 | 0.020 |
| GSE63496 | SRR1658025 | RNA-seq | 136 | 3601 | 0.020 |
| GSE62047 | SRR1596500 | GRO-seq | 10496 | 236794 | 0.849 |
| GSE62047 | SRR1596501 | GRO-seq | 2926 | 44319 | 0.141 |
| GSE71901 | SRR2153508 | GRO-seq | 104293 | 4720754 | 5.098 |
| GSE71901 | SRR2153509 | GRO-seq | 109035 | 4866057 | 5.608 |
| GSE71901 | SRR2153510 | GRO-seq | 40897 | 4175377 | 4.167 |
| GSE41357 | SRR580356 | HITS-CLIP | 4305 | 514864 | 3.285 |
| GSE41357 | SRR580357 | HITS-CLIP | 2239 | 366095 | 2.898 |
| GSE42701 | SRR628446 | HITS-CLIP | 3241 | 69343 | 0.802 |

### RT mispriming occurs in multiple RNA-seq based technologies and leads to misinterpretation of data

In addition to short RNA-seq experiments, several technologies utilize 3’ adapter ligation followed by reverse transcription for library preparation (Fig. 1A) including HITS-CLIP, NET-seq, GRO-seq, total/mRNA-seq, RIP-seq and RIBO-seq. Many of these technologies rely on specific positions of cDNA peaks to identify the binding sites of RNA binding proteins (CLIP based approaches), or RNA polymerase (GRO-seq and NET-seq) or ribosome footprinting sites (RIBO-seq). Since RT mispriming produces false cDNA peaks, we hypothesized that mispriming could lead to misinterpretation of data from sequencing technologies that rely on the specific position of cDNA peaks [6,9,10,16]. In HITS-CLIP, RT mispriming was recently shown to produce false peaks identified by looking for sequences matching the first 6-7 bases of the 3’ adapter in a 200 bp region spanning the cDNA peaks [7]. This approach would fail to identify mispriming sites that have scattered matches to the 3’ adapter beyond the first 2 bases. Using our pipeline, we were able to detect and filter out several mispriming sites in two independent published CLIP datasets (Table 2, Fig. 4A-C) [17,18]. In addition to mispriming sites with 6-7 base matches, we identified mispriming sites with scattered matches to the 3’ adapter that would be missed by the existing pipelines (Figure S4A, B). In order to check if RT mispriming could lead to false positive peak calls, we compared the number of peaks identified using the peak caller pyicoclip in one of the downloaded CLIP datasets before and after filtering for misprimed reads [19]. We detected ~2.5% peaks as false positive peak calls that could be misinterpreted as binding sites.

**Figure 4.**
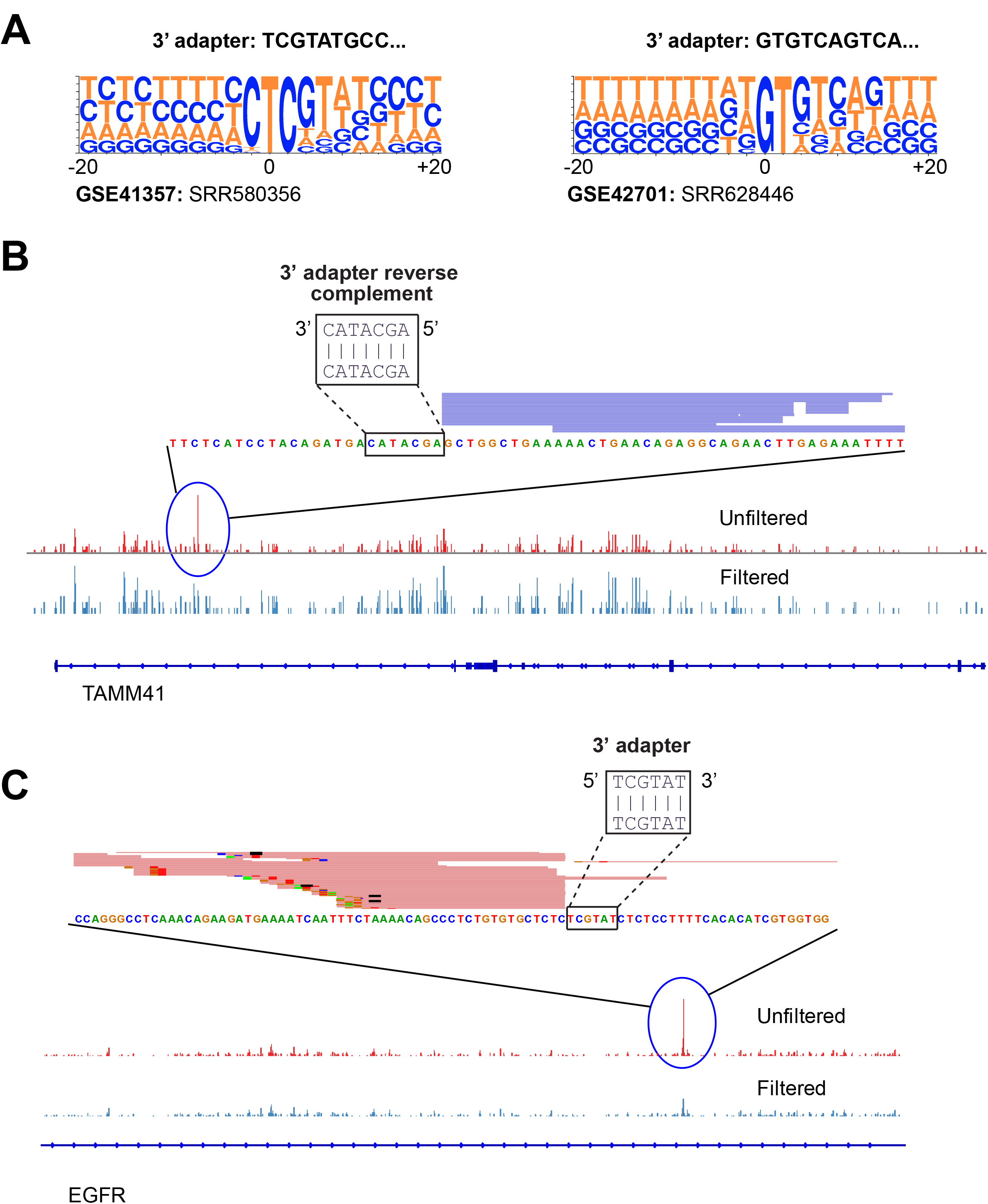
(A) Sequence logos showing sequence enrichment spanning 20 bp around mispriming sites identified by our pipeline for two published HITS-CLIP datasets. (B, C) View of a HITS-CLIP peak adjacent to a sequence matching the 3’ adapter.

**Table 2:** RT mispriming events identified by our pipeline in mRNA-seq and short RNA-seq datasets we generated.

| Sample | Technique | Number of mispriming sites | Number of misprimed reads | Percent of reads misprimed |
| --- | --- | --- | --- | --- |
| T98G EZH2 IP BR1 | Short RNA-seq | 36558 | 1531591 | 44.070 |
| T98G EZH2 IP BR2 | Short RNA-seq | 37203 | 1704240 | 28.926 |
| T98G IgG IP BR1 | Short RNA-seq | 10439 | 227480 | 2.766 |
| T98G IgG IP BR2 | Short RNA-seq | 12968 | 302368 | 3.808 |
| T98G Input BR1 | Short RNA-seq | 1318 | 32800 | 0.267 |
| T98G Input BR2 | Short RNA-seq | 3009 | 91975 | 0.514 |
| U87MG EZH2 IP BR1 | Short RNA-seq | 1878 | 49135 | 4.870 |
| U87MG EZH2 IP BR2 | Short RNA-seq | 46293 | 2380088 | 54.748 |
| U87MG IgG IP BR1 | Short RNA-seq | 591 | 11015 | 1.040 |
| U87MG IgG IP BR2 | Short RNA-seq | 19981 | 535942 | 8.182 |
| U87MG Input BR1 | Short RNA-seq | 67 | 1770 | 0.069 |
| U87MG Input BR2 | Short RNA-seq | 1920 | 57762 | 0.593 |
| U87MG Total RNA-seq BR1 | RNA-seq | 2890 | 37395 | 0.066 |
| U87MG Total RNA-seq BR2 | RNA-seq | 3638 | 45185 | 0.064 |

We next applied our pipeline to identify mispriming events in GRO-seq datasets, another technology that relies on the specific position of cDNA peaks. GRO-seq is primarily used to identify sites of elongating RNA polymerase based on the position of cDNA peaks. High density of cDNA reads close to a gene’s transcription start site relative to the gene body indicates RNA polymerase promoter-proximal pausing. The extent of pausing is measured in terms of the pausing index (PI), calculated as the ratio of the number of reads per kilobase mapping close to transcription start sites (within 300 bp spanning the TSS) to reads mapping to the gene body (TSS + 250 bp to TSS +2250 bp) [20–22]. In several GRO-seq datasets we were able to identify 10,000 - 50,000 mispriming sites accounting for millions of reads in some datasets (Table 2, Fig. 5A-B and Figure S5) [23,24]. Since GRO-seq utilizes the position-specific count of cDNA reads to detect RNA polymerase pausing, we suspected that artefactual fluctuations in read densities as a result of mispriming could lead to mis-identification of pause sites. To test for this, we identified and filtered out misprimed reads from a published GRO-seq dataset and analyzed it for differences in pausing index before and after filtering. Differences in PI between unfiltered and filtered data would highlight cases of erroneous measurements of RNA polymerase pausing. For one of the GRO-seq datasets (GSE71898: SRR2153508), we found 230 protein-coding genes where the unfiltered dataset showed at least 2-fold difference in PI values compared to the filtered dataset (Fig. 5C, D) [24]. These protein-coding genes include cases where mispriming leads to higher PI (indicating promoter-proximal pausing) and some that show lower PI (indicating higher elongation rates).

**Figure 5.**
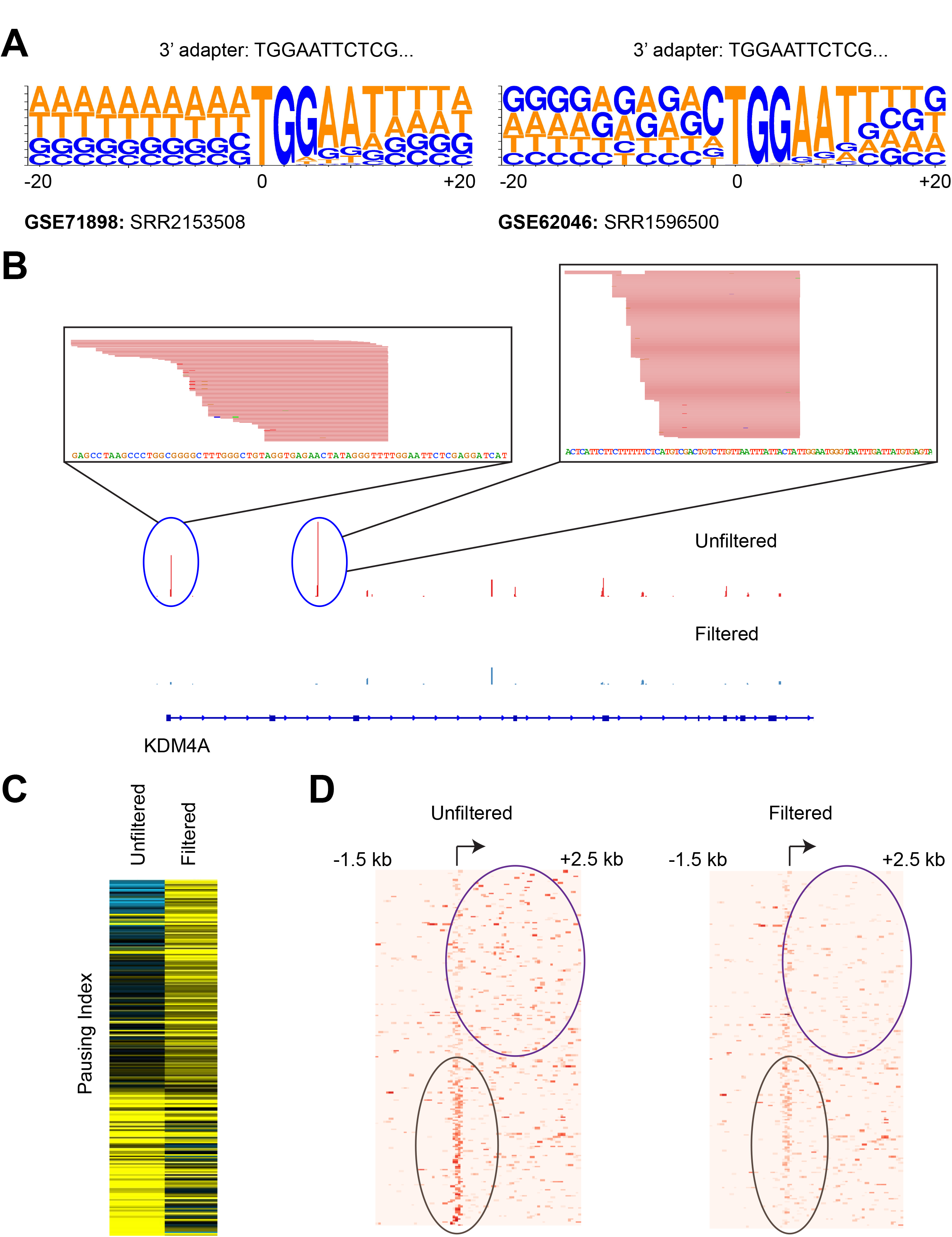
(A) Sequence logos showing sequence enrichment spanning 20 bp around mispriming sites identified by our pipeline for two published GRO-seq datasets. (B) View of GRO-seq peaks adjacent to sequence matching the 3’ adapter sequence. (C) Heatmap showing pausing index (PI) for 230 genes that show at least 2-fold difference in PI before and after filtering for mispriming reads. Rows are ordered by decreasing ratio of PI in filtered to PI in unfiltered dataset. (D) Heatmap showing distribution of reads proximal to transcription start site and gene body in unfiltered (left) and filtered dataset (right) for 230 genes shown in C.

### RT mispriming can be avoided by using TGIRT-seq

One way to address sequencing artifacts arising from RT mispriming, as proposed for mammalian NET-seq and HITS-CLIP, is to include a degenerate barcode on the 5’ end of the 3’ adapter sequence [6,7]. With this approach, misprimed reads can be removed by collapsing reads with identical 3’ adapter barcode sequences. Although this approach helps with filtering out mispriming-induced artifacts from downstream analysis, there would be a substantial loss of sequencing reads. As an alternative, we propose RNA-seq using thermostable group II intron-encoded reverse transcriptase (TGIRT-seq) to avoid RT mispriming. TGIRT-seq is a relatively new RNA-seq workflow that utilizes template switching to link 3’ adapter sequences to the synthesized cDNA. In contrast to other small RNA library preparation methods (Fig. 1A, right), TGIRT-seq synthesizes cDNA by template switching from a pre-annealed 3’ adapter RNA/DNA heteroduplex, skipping the priming step (Fig. 6A) [11,12]. Since this approach does not involve cDNA synthesis dependent on specific priming by RT-primer, we hypothesized that artifacts from RT mispriming would not occur in data generated using TGIRT-seq. To test this, we performed short RNA sequencing of immunoprecipitated RNA from non-specific control IgG using TGIRT-seq. By performing TGIRT-seq on the control IP sample, we were able to compare short RNA peaks from coding exons with background peaks from other classes of short RNAs (miRNAs, snoRNAs etc.,) as an internal control. Although we found several thousand reads mapping to multiple classes of short RNAs, we were unable to detect short RNA peaks from coding exons we previously observed with the NEB small RNA library preparation kit (Fig. 6B). This shows that TGIRT-seq can help avoid RT mispriming without compromising on read coverage at transcripts that exist in the cell.

**Figure 6.**
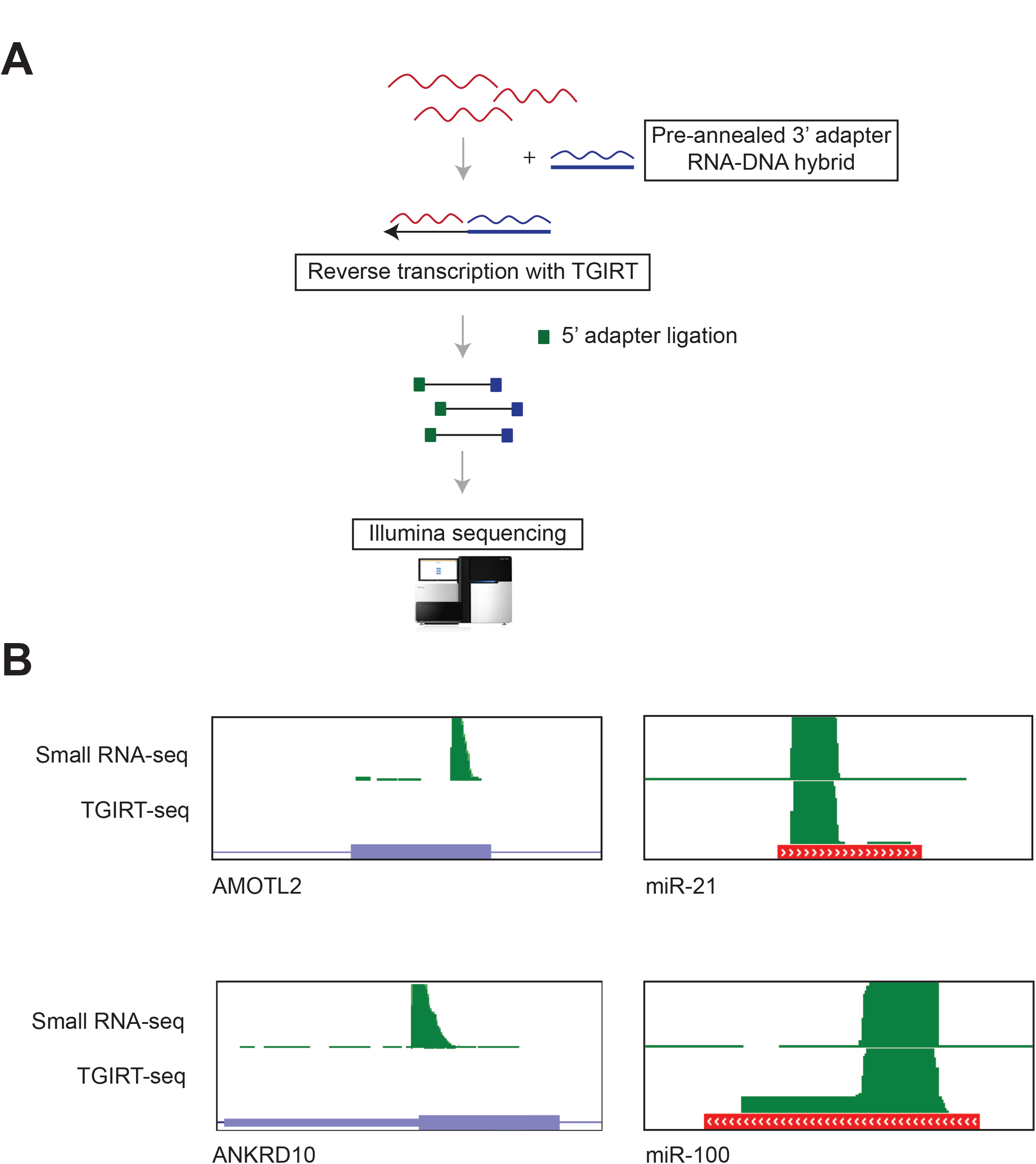
(A) Overview of TGIRT-seq protocol. 3’ adapter RNA-DNA hybrid is annealed and used in a template switching reaction with TGIRT to synthesize cDNA molecules linked to the 3’ adapter. This is followed by ligation of 5’ adapter, PCR amplification and sequencing. (B) Genome browser views of misprimed exonic cDNA peaks (left) and miRNA peaks (right) in a short RNA-seq library prepared with NEB small RNA library kit and TGIRT-seq.

## Discussion

RNA-seq based technologies are widely used to answer questions relevant to gene expression changes, protein-RNA interactions, RNA-RNA interactions, identifying RNA secondary structure and RNA polymerase dynamics during transcription [6,9,25–28]. The quality and accuracy of sequencing data relies on efficient priming by the RT-primer during cDNA synthesis, ligation of adapters and priming by primers during PCR amplification. Although sequencing bias during PCR amplification and adapter ligations have been previously addressed and solutions proposed, bias during reverse transcription is underappreciated [5,29]. Data from RNA-seq technologies can be contaminated with incorrect cDNA ends as a result of mispriming by the primer used for reverse transcription (RT primer). If unaccounted for during analysis, misprimed reads can lead to misinterpretation of data. Here we developed an analysis pipeline aimed at identifying and eliminating misprimed reads from aligned RNA-seq datasets. An earlier approach to identifying mispriming events was based on looking for cDNA pile-ups adjacent to sequences matching the first 6-7 bases of the 3’ adapter (complementary to the RT primer). Based on detailed analysis of short RNA-seq datasets, we found that the RT-primer does not necessarily require base-pairing over a stretch of 6-7 bases to cause mispriming but instead can occur at sites with a match to only the first two bases followed by scattered complementarity. Thus, the earlier approach relying on a 6-7 base match is insufficient to identify all mispriming sites. We applied our mispriming identification pipeline to filter out mispriming artifacts in several published datasets from multiple technologies including short RNA-seq, total RNA-seq, HITS-CLIP and GRO-seq. We further show how failure to remove misprimed reads could lead to misinterpretation of data.

In our short RNA-seq datasets, we found reads mapping to coding exons in addition to known classes of short RNAs (miRNAs, snoRNA etc.). These cDNA read pile-ups at coding exons showed flush 3’ ends that ended exactly before sequence matching the 3’ adapter sequence that we had used to generate cDNA libraries. This is a key characteristic of a mispriming event that were previously shown for HITS-CLIP and NET-seq [6,7]. In case of both HITS-CLIP and NET-seq, mispriming artifacts were previously detected by finding sequences spanning a region around cDNA peaks that matched the first 6-7 bases of the 3’ adapter (complementary to the RT primer). We however were able to detect several cDNA peaks with only scattered complementarity beyond the first 2 bases of RT-primer. This suggests that the mispriming sites previously identified for HITS-CLIP and NET-seq represent only a subset of all mispriming sites.

Using our pipeline we were able to identify thousands of mispriming sites in multiple short RNA-seq datasets. We found that the number of mispriming sites were significantly lower for input libraries (short RNAs from total RNA pool) compared to immunoprecipitated samples. This is likely attributable to relatively smaller pool of distinct RNA molecules in the immunoprecipitated sample. We next applied this pipeline to several published datasets from several technologies including short RNA-seq, total/mRNA-seq, HITSCLIP and GRO-seq. We were able to detect hundreds to thousands of mispriming sites from multiple datasets. RNA-seq is primarily performed to detect differential expression of genes and splicing changes across transcripts between two conditions. In both kinds of analysis, results are based on read coverage across a large region of a gene or transcript. Even though we were able to detect hundreds of mispriming events in multiple mRNA-seq datasets, they had negligible impact on gene expression and splicing (data not shown) [30,31]. On the contrary, with technologies that rely on specific positions of cDNA peaks (HITS-CLIP and GRO-seq), the mispriming events that we identified showed a much greater impact on data interpretation. In case of HITS-CLIP, we identified thousands of peaks that would be misidentified as binding sites and for GRO-seq we identified several genes where mispriming could lead to misinterpretation of RNA polymerase elongation dynamics. This shows that RT mispriming affects multiple RNA-seq datasets and can lead to widespread misinterpretation of data.

One proposed experimental approach to avoid RT mispriming is to alter the 5’ end of the 3’ adapter sequence to contain degenerate barcodes that can later be used to collapse reads with identical 3’ adapter sequence. Although this approach helps remove misprimed reads from downstream analysis, this can lead to loss of sequencing data. As an alternative, we propose the use of TGIRT-seq that lacks RT-priming step during library preparation. This approach completely eliminates mispriming artifacts from the library (Fig. 6B).

In this manuscript, we provide evidence for how RT mispriming contaminates and leads to misinterpretation of data for several RNA-seq libraries across multiple technologies. As a solution we provide an analysis pipeline to filter out misprimed reads from sequencing data that will be useful for future analysis utilizing published datasets. We also provide an alternative experimental approach to avoid RT mispriming during RNA-seq library preparation.

## Materials and Methods

### Cell lines and reagents

T98G and U87MG (ATCC-CRL-1690 and ATCC-HTB14) were grown in EMEM with 10% FBS. All cell lines were maintained at 37°C and 5% CO_2_. Antibodies used were: EZH2 (Active Motif 39875) and IgG2a (Sigma Aldrich M5409).

### Nuclear lysis

All RIP-seq experiments were performed using nuclear lysates. Cells were incubated in hypotonic solution (10 mM KCL, 10 mM HEPES pH 7.5, 1.5 mM MgCl_2_ and 2 mM DTT) for 5 min and spun down by centrifugation. 1 mg/ml digitonin (Sigma Aldrich D141) was then added to the lysate resuspension and further incubated for 10 min with constant mixing. Cells were then mechanically lysed using a dounce homogenizer (15 times) and spun down. Pelleted cells were then resuspended in NP40 lysis buffer (150 mM KCL, 50 mM HEPES pH 7.5, 5 mM EDTA, 0.5 % IGEPAL and 2 mM DTT) and subjected to mild sonication (6 cycles, 10 sec on and 30 sec off). All the steps in this procedure were performed at 4°C.

### RNA-seq and RIP-seq

RNA-seq experiments were performed on poly-A selected mRNA (Bioo 512980) as previously described [31]. RNA immunoprecipitations were performed on nuclear lysates from 4 × 150 cm^2^ plates of cells using 10 μg of specific antibody per experiment. Magnetic bead preparation and immunoprecipitations (IP) were performed as previously described [31]. 10% of the nuclear lysate was separated as input for total RNA-seq. Following IP, beads were resuspended in Trizol (Thermo Fisher Scientific 15596026) and RNA was extracted as per manufacturer’s instructions. Small RNA RIP-seq libraries were prepared using NEB-next small RNA library preparation kit (NEB E7330) for IP and input. Following library preparation, 20-50 bases long products were size-selected from 6% TBE gel as per manufacturer’s instructions.

### Identification of EZH2-enriched exonic cDNA peaks

Small RNA RIP-seq reads were mapped to the human genome (UCSC version hg19) using BWA and reads mapping to exons were counted using bedtools [32,33]. For GBM cells where RIP-seq experiments were performed in replicates, reads mapping to exons were compared to negative controls (IgG and input) using DESeq2 to identify exons significantly enriched in the EZH2 short RNA RIP [34]. To be identified as an EZH2-enriched coding exon, the number of normalized reads mapping to exons were required to be significantly higher (FDR-corrected *P* < 0.05) by at least 2-fold in the EZH2 RIP in comparison to both negative controls. In addition, exonic cDNA peaks were called using pyicoclip, and only peaks containing EZH2-enriched coding exon were included [19].

### Analysis of published datasets

All external datasets listed in Table 2 were downloaded from GEO. Adapters from Fastq files were removed using cutadapt and mapped to the human genome (UCSC version hg19) using BWA [35,36]. Mispriming sites were identified from aligned files using our scripts (Mispriming_finder.py and Filter_misprimes.py) and misprimed reads were removed using bedtools. Scripts used for this analysis are publically available on github (https://github.com/haridh/RT-mispriming). For HITS-CLIP, peaks were called using pyicoclip before and after filtering out misprimed reads. For GRO-seq, pausing index at genes containing misprimed reads was compared before and after filtering out misprimed reads. Pausing index was calculated as the ratio of the number of reads per kilobase mapping close to transcription start sites (within 300 bp spanning TSS) and gene body (TSS + 250 bp to TSS +2250 bp).

## Acknowledgments

We thank Nathan S. Abell for optimizing the nuclear lysis protocol, Anna Battenhouse for assistance with aligning sequencing data, Alan Lambowitz and Ryan Nottingham for providing TGIRT-seq reagents and helpful discussion regarding the protocol, Robert Darnell and Christopher Park for helpful discussions about RT mispriming, the Genomic Sequencing and Analysis Facility at UT Austin and MD Anderson Cancer Center-Science Park NGS Facility for Illumina sequencing. The Science Park NGS Facility was supported by CPRIT Core Facility Support Grant RP120348. We also thank the Texas Advanced Computing Center (TACC) at UT Austin for the use of computational facilities. This work was funded in part by grants from the Cancer Prevention and Research Institute of Texas (RP120194) and the National Institutes of Health (NIH) (CA198648) to V.R.I. None of the authors have any competing interests.

## Data

Primary sequencing data generated in this study is available at NCBI’s GEO database (www.ncbi.nlm.nih.gov/geo/query/acc.cgi?acc=GSE85163). Python scripts used to identify RT mispriming events are available on github (https://github.com/haridh/RT-mispriming).

## Supplementary Figures

**Figure S1.**
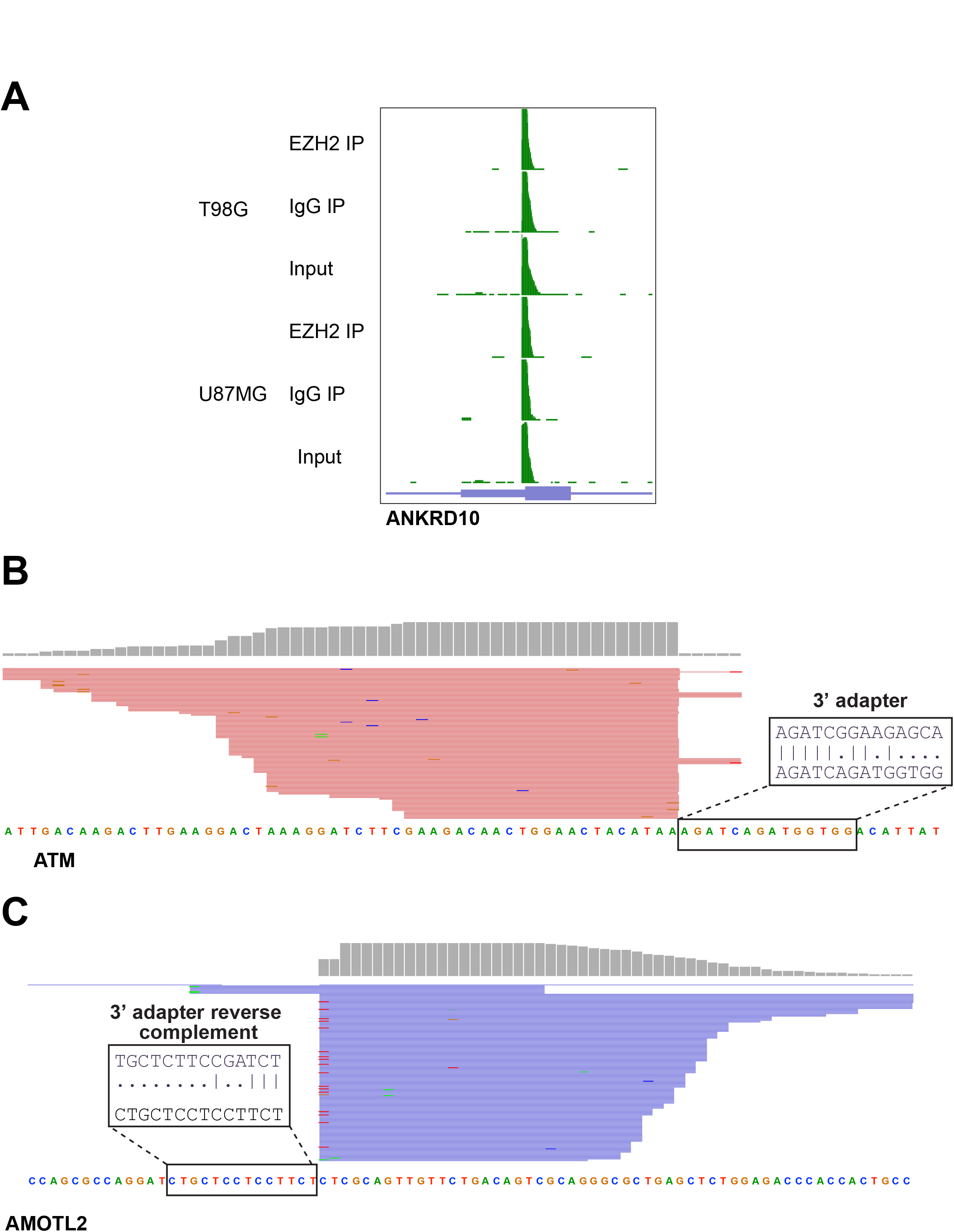
(A) Genome browser view showing cDNA peak enriched at a specific exon in EZH2 short RNA RIP but also present in IgG IP and input controls across different cell lines. (B) View of a cDNA peak from short RNA-seq showing 3’ flush ends adjacent to a sequence similar to the 3’ adapter (first 5 bases matching). (C) View of a cDNA peak adjacent to a sequence with first 3 bases followed by scattered matches to the 3’ adapter.

**Figure S2.**
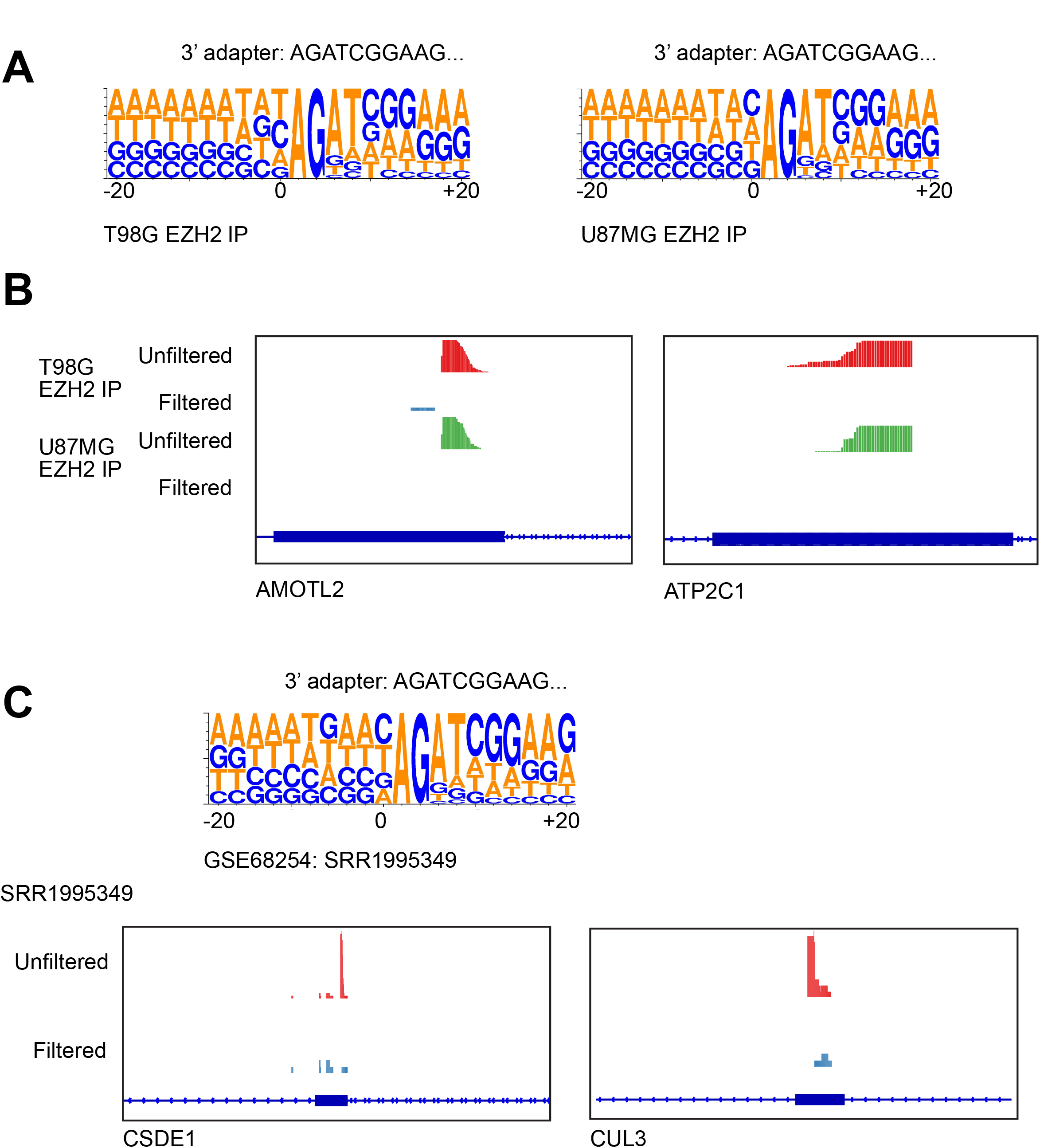
(A) and (C, Top) Sequence logos showing sequence enrichment spanning 20 bp around mispriming sites identified by our pipeline for short RNA-seq datasets we generated (A) and a published short RNA-seq dataset (C, top). (B) and (C, bottom) View of cDNA peaks in EZH2 IP samples before and after filtering out mispriming events identified by our pipeline for short RNA-seq dataset we generated (B) and published short RNA-seq dataset (C, bottom).

**Figure S3.**
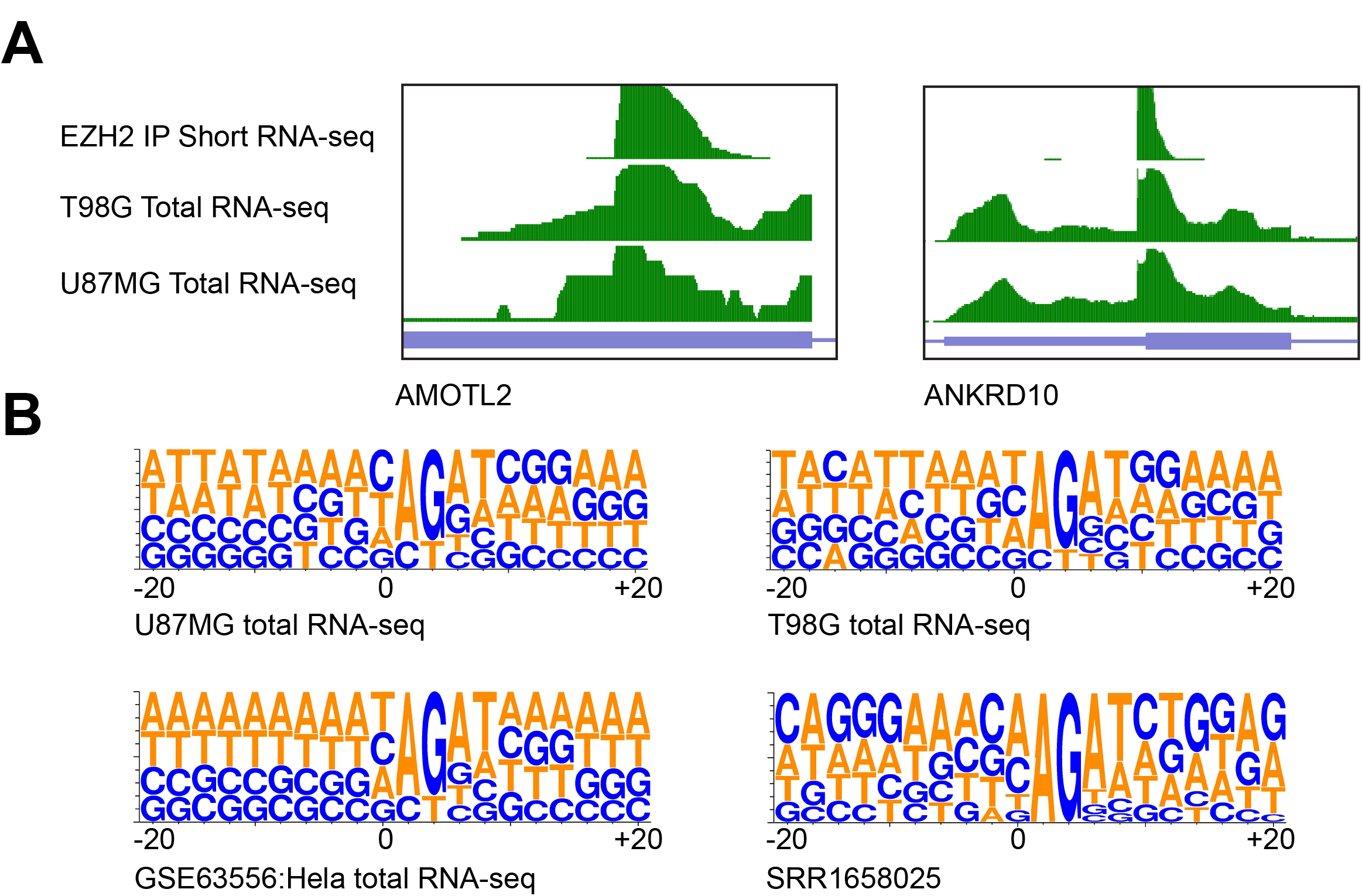
(A) Genome browser views of misprimed peaks in total RNA-seq datasets that coincide with misprimed peaks in short RNA-seq datasets. (B) Sequence logos showing sequence enrichment spanning 20 bp around mispriming sites identified by our pipeline for total RNA-seq datasets we generated and two datasets published previously.

**Figure S4.**
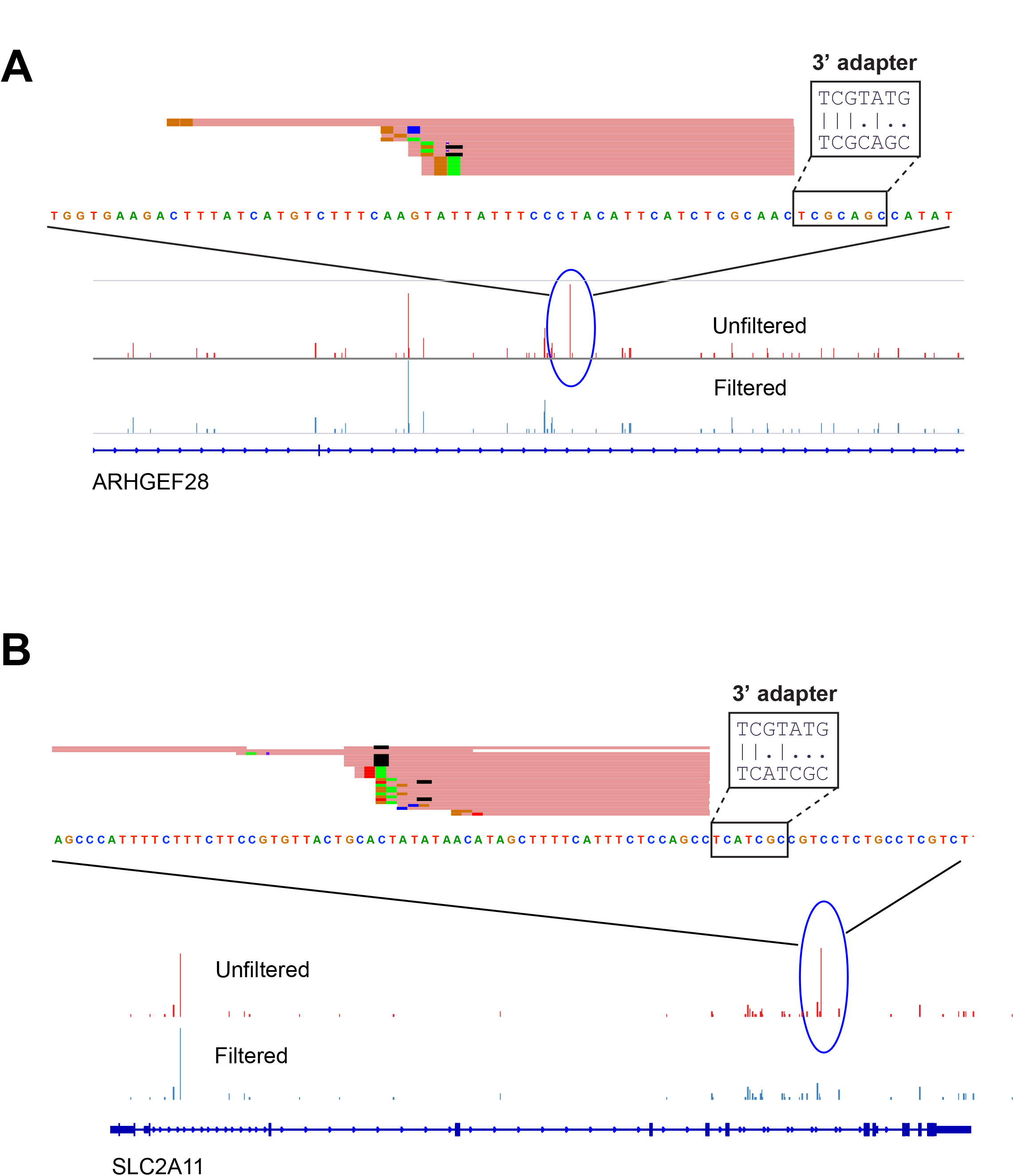
(A) and (B), View of a HITS-CLIP peak adjacent to a sequence with scattered matches to the 3’ adapter sequence.

**Figure S5.**
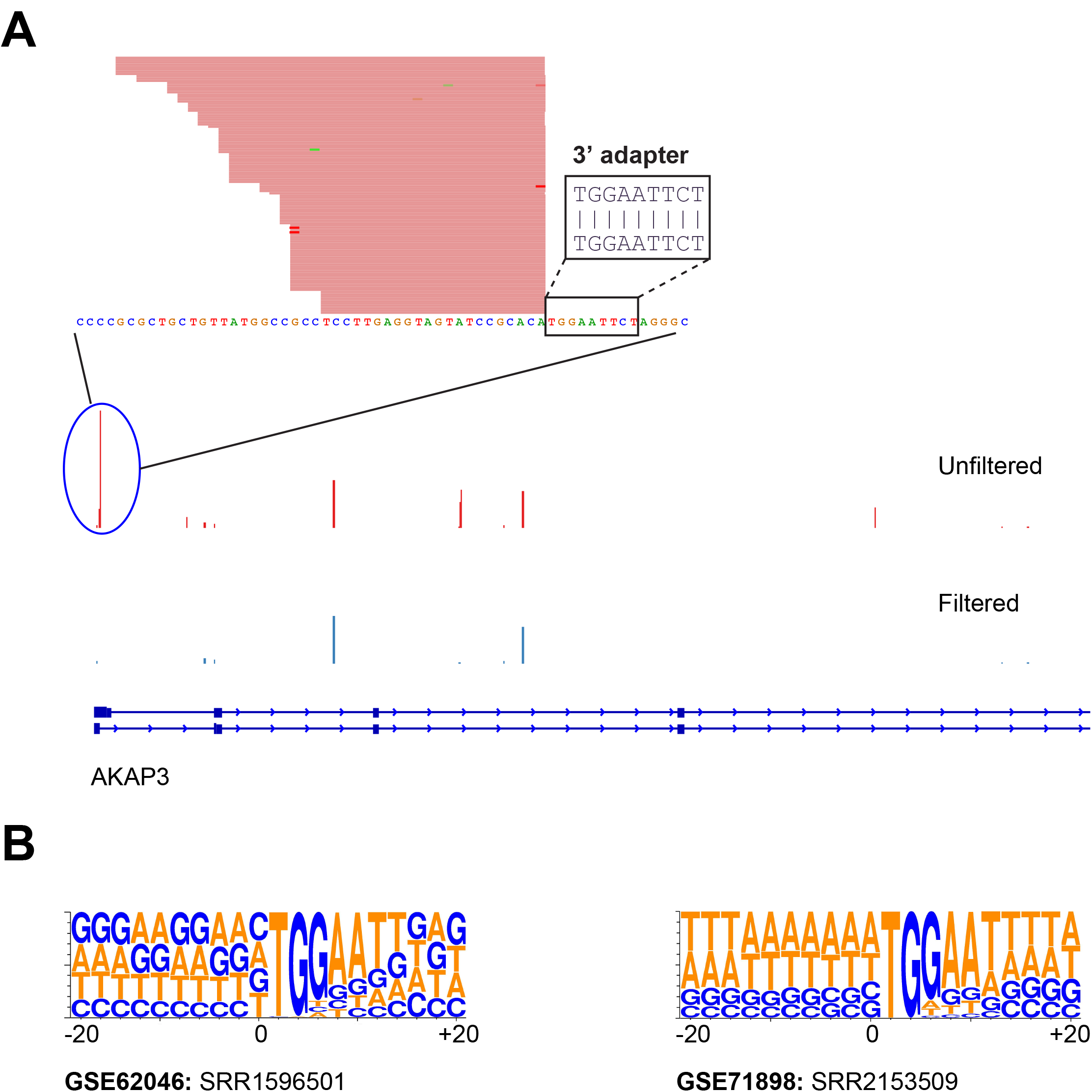
(A) View of a GRO-seq peak adjacent to a sequence matching the 3’ adapter sequence. (B) Sequence logo showing sequence enrichment spanning 20 bp around mispriming sites identified by our pipeline for two other published GRO-seq datasets.

